# High-throughput miRNA-sequencing of the human placenta: expression throughout gestation

**DOI:** 10.1101/2021.02.04.429392

**Authors:** Tania L Gonzalez, Laura E Eisman, Nikhil V Joshi, Amy E Flowers, Di Wu, Yizhou Wang, Chintda Santiskulvong, Jie Tang, Rae A Buttle, Erica Sauro, Ekaterina L Clark, Rosemarie DiPentino, Caroline A Jefferies, Jessica L Chan, Yayu Lin, Yazhen Zhu, Yalda Afshar, Hsian-Rong Tseng, Kent Taylor, John Williams, Margareta D Pisarska

## Abstract

**Background:** Altered placenta miRNA abundance may impact the maternal-fetal interface and pregnancy outcomes. Understanding miRNA changes across gestation is essential before miRNAs can be used as biomarkers or prognostic indicators during pregnancy.

**Materials & Methods:** Using next-generation sequencing, we characterize the normative human placenta miRNA transcriptome in first (N=113) and third trimester (N=47).

**Results:** There are 801 miRNAs expressed in both first and third trimester, including 182 with similar expression across gestation (P≥0.05) and 182 significantly different (FDR<0.05). Of placenta-specific miRNA clusters, C14MC is more upregulated in first trimester and C19MC is more highly expressed overall.

**Conclusion:** This work provides a rich atlas of healthy pregnancies to direct functional studies investigating the epigenetic differences in first and third trimester placentae.

**Lay Abstract:** The human body produces microRNAs which affect the expression of genes and proteins. This study uses next generation sequencing to identify the microRNA profile of first and third trimester human placentae using a large cohort (N=113 first, N=47 third trimester). All pregnancies resulted in healthy babies. We identify microRNAs with significantly different expression between first and third trimester, as well as stably expressed microRNAs. This work provides a baseline for future studies which may use microRNAs to monitor maternal-fetal health throughout pregnancy.

## Introduction

The placenta plays a critical role in fetal development, forming the interface between the developing fetus and mother. It has multiple functions including provision of oxygen and nutrients to the fetus, removal of waste products, and constitutes an important barrier protecting the fetus from pathogens and environmental toxins throughout gestation. The formation and development of the placenta (placentation) that begins upon implantation and continues throughout the first trimester of pregnancy lays the foundation for a placenta leading to a healthy gestation. After implantation, the placenta produces growth factors, cytokines and hormones which target maternal physiological systems, facilitating the provision of additional blood flow and nutrient delivery to the fetus.^1^ During the first trimester, cytotrophoblast cells differentiate into extravillous trophoblasts and syncytiotrophoblasts. Extravillous trophoblasts migrate by invading and remodeling the maternal decidual extracellular matrix and spiral arteries.^2–5^ This occurs during a relative state of hypoxia, which promotes trophoblast invasion and angiogenesis.^6–9^ It is also during this time that cytotrophoblasts fuse into syncytiotrophoblasts, creating a multinucleated epithelium that lines the intervillous space and also produces hormones, including estrogen and progesterone for pregnancy maintenance and lactogen for fetal metabolism, growth and development.^10–12^ The complexities of normal placental development are under the control of various epigenetic modifications that may be altered, ultimately leading to abnormal placentation and adverse outcomes.^13,14^

MicroRNAs (miRNAs) are non-coding, single-stranded RNA molecules of about 22 nucleotides in length.^15^ They are important post-transcriptional regulators of gene expression. They bind to RNA transcripts, causing RNA cleavage or messenger RNA translational repression, regulating 30-50% of all mammalian protein-coding genes.^16,17^ While some miRNAs are universally expressed, others are expressed preferentially or exclusively in certain tissues.^18^ Two large miRNA clusters are enriched in placenta, the chromosome 14 miRNA cluster (C14MC) and the chromosome 19 miRNA cluster (C19MC).^19,20^ C14MC is a large, imprinted, maternally-expressed miRNA cluster, with several members predominantly expressed in placenta and epithelial tissues.^18^ C19MC is a large, imprinted, paternally-expressed miRNA cluster whose members have highest expression in placenta and cancer, with relatively weak expression in other tissue.^18,20–24^

Widespread regulation of gene expression by miRNAs, the presence of placental-specific miRNAs, and miRNA expression differences in various trophoblastic cell lines^25^ suggests a role for miRNAs in trophoblast behavior and placental development and function. Furthermore, altered miRNA expression in the placenta may be involved in abnormal placentation and related pregnancy-associated diseases, including preeclampsia^26–40^ and intrauterine growth restriction (IUGR).^39–43^ Given that placental function changes from the first trimester during a critical state of placental development and continues to function for appropriate fetal development, epigenetic regulation through miRNAs likely play a key role. In addition, due to their small size and stability, miRNAs are potential biomarkers of disease, particularly miRNAs from plasma exosomes,^44–46^ however, it is essential to know how their expression varies across gestation. Previous studies comparing miRNA expression in first and third trimester placentae are limited to microarray analyses and have had conflicting results.^25,47,48^ We performed next-generation sequencing (NGS) and expression analysis to identify and compare the normative miRNA signatures in first and third trimester placentae of healthy pregnancies resulting in delivery.

## Materials and Methods

### Study Population

The study population consisted of 157 singleton pregnancies with available first trimester placental tissue (N=110), third trimester placental tissue (N=44) or both (N=3 in both groups), obtained between 2009 and 2018. Mothers with pre-existing diabetes or hypertension were excluded. All subjects were enrolled under IRB approved protocols (Pro00006806, Pro00008600). All pregnancies had a normal karyotype and resulted in the delivery of a viable infant.

### Analysis of demographic data

Demographic data included parental ages, races and ethnicities, maternal pre-pregnancy body mass index, gestational age at chorionic villus sampling (CVS), fetal sex, maternal medical history and medication use, pregnancy complications, mode of delivery, gestational age at delivery, and birth weight. Means and standard deviations were reported for continuous variables. Proportions were reported as percentages. Demographics were compared between patients in the first and third trimester placenta, excluding 3 subjects sequenced in both groups to eliminate duplicate information. T-test was used for normally distributed continuous variables, and the Wilcoxon rank-sum test was used for non-parametric data. Fisher’s exact test was used when appropriate. For comparison of categorical variables, the chi-square test was used.

### Collection of first trimester placental samples

Samples from the first trimester of pregnancy were collected between 70-102 days gestation during CVS procedures done for prenatal diagnosis. Samples used for research consisted of extra tissue which is normally discarded after sending chorionic villi specimens for prenatal genetic diagnostic testing. Fetal-derived chorionic villi (first trimester placenta tissue) were cleaned and separated from any maternally-derived decidua (non-placenta tissue). Tissue samples (5-25 mg) were kept on ice and submerged in RNA*later* RNA stabilization reagent (QIAGEN, Hilden, Germany) within 30 minutes of collection and stored at −80°C.

### Collection of third trimester placental samples

Samples from the third trimester of pregnancy were collected between 254-290 days gestation, after delivery of a viable neonate. Samples used for research consisted of tissue which would have otherwise been discarded. One centimeter cubed of placental tissue samples were obtained immediately after delivery from the fetal side of the placenta near the site of cord insertion beneath the amnion. The samples were cleaned and submerged in RNA*later* RNA stabilization reagent (QIAGEN) and stored at −80°C.

### RNA extraction from first trimester placenta

RNA was extracted from leftover CVS tissue utilizing a method optimized for delicate tissue.^49,50^ Briefly, tissue samples were thawed on ice with 600 μl of RLT Plus lysis buffer (QIAGEN) containing 1% β-mercaptoethanol. Tissue was homogenized by passing at least 10 times through progressively thinner gauge needles (22G, 25G, 27G) attached to an RNase-free syringe. Homogenates were loaded onto AllPrep spin columns and the remainder of sample processing was performed following manufacturer instructions using the AllPrep DNA/RNA/miRNA Universal Kit (QIAGEN). RNA was eluted with 30-45 μl of RNase-free water at room temperature and the elution was passed through the column twice to improve yields, as previously described.^49,50^ The average RNA integrity number (RIN) for sequenced first trimester placenta samples was 8.87.

### RNA extraction from third trimester placenta

Third trimester placenta tissue was thawed on ice, then a quarter of collected tissue was diced with RNase-free blades coated in RNA*later* buffer, tissue was sonicated on ice in lysis buffer (600 μL RLT Plus lysis buffer with 1% β-mercaptoethanol) using 5 second pulses on a low setting (#2) until tissue fragments were small enough to complete homogenization with RNase-free needles. Further extraction was performed as described for first trimester tissue. The average RIN for sequenced third trimester placentae was 8.84.

### Library preparation and miRNA sequencing

A miRNA sequencing library was prepared from total RNA using the QIASeq miRNA Library Kit (QIAGEN, Hilden, Germany). A pre-adenylated DNA adapter was ligated to the 3’ ends of miRNAs, followed by ligation of an RNA adapter to the 5’ end. A reverse-transcription primer containing an integrated Unique Molecular Index (UMI) was used to convert the 3’/5’ ligated miRNAs into cDNA. After cDNA cleanup, indexed sequencing libraries were generated via sample indexing during library amplification, followed by library cleanup. Libraries were sequenced on a NextSeq 500 (Illumina, San Diego, CA) with a 1×75 bp read length and an average sequencing depth of 10.64 million reads per sample.

### Differential expression analysis of miRNAs

The demultiplexed raw reads were uploaded to GeneGlobe Data Analysis Center (QIAGEN) at https://www.qiagen.com/us/resources/geneglobe/ for quality control, alignment and expression quantification. Briefly, 3’ adapter and low quality bases were trimmed off from reads first using *cutadapt* v1.13 with default settings,^51^ then reads with less than 16bp insert sequences or with less than 10bp UMI sequences were discarded. The remaining reads were collapsed to UMI counts and aligned sequentially to miRBase release 21 mature and hairpin RNA databases using *Bowtie* v1.2.^52,53^ The UMI counts of each miRNA category were quantified, and then normalized by a size factor-based method in the R package *DESeq2* v1.22.2 (Bioconductor).^54^ Data were averaged across all samples in each group (first and third trimester) for each respective miRNA and were reported as baseMean. The R package *FactoMineR* v1.41 was used to conduct principal components analysis (PCA), which was used to investigate clustering and potential outliers. Differential expression analysis was performed with *DESeq2* to compare first versus third trimester expression, adjusting for fetal sex. Each miRNA was fitted into a negative binomial generalized linear model, and the Wald test was applied to assess the differential expressions between two sample groups (P value). The Benjamini and Hochberg procedure was applied to adjust for multiple hypothesis testing, and those with false discovery rate (FDR) less than 0.05 were selected as significantly differentially expressed miRNAs. The genome locations of miRNAs were identified by cross-referencing mature miRNA IDs with precursor miRNA accession IDs in miRBase release 21, then using R package *biomaRt* v2.45.8 and Ensembl release 91 (which contains miRBase release 21) to retrieve chromosome locations.^55,56^ For miRNAs derived from more than one precursor, all chromosomal locations were counted in bar plots (e.g. hsa-miR-1184 is encoded by three precursors on chromosome X, thus was counted three times) and each miRNA was plotted once per chromosome in scatter plots of genome distribution.

### Analysis of miRNA expression

Counts normalized for sequencing depth (baseMeans) were used as a measure of expression since miRNA lengths do not vary substantially. All expressed miRNAs were defined as any with baseMean>10 in both first or third trimester placenta groups. The miRNAs common in both first and third trimester placentae, also known as similarly expressed, were defined by P≥0.05, absolute fold-change≤2, and baseMean>10 in both trimesters. The P≥0.05 threshold excludes all significantly different miRNAs (FDR<0.05) as well as other potentially different miRNAs (P<0.05). Differentially expressed miRNAs were defined as FDR<0.05, absolute fold-change>2, and baseMean>10 in both trimesters. Higher expression thresholds were selected for target enrichment analysis when needed (next section). For miRNAs in C14MC and C19MC, these filters were applied: baseMean>1 and P≥0.05 (similarly expressed) or baseMean>1 and FDR<0.05 (differentially expressed).

### Enrichment analysis of predicted miRNA target genes

Ingenuity Pathways Analysis (IPA) software’s microRNA Target Filter application (QIAGEN, Redwood City, CA, USA, http://www.qiagenbioinformatics.com/IPA) was used to generate a list of target RNAs based on sequence and experimental confirmation. Targets were included if biochemically confirmed using human tissue or non-species methods (sourced from QIAGEN’s curated Ingenuity Knowledge Base, or the publicly available miRecords or TarBase). IPA’s Core Analysis function was used to test the hypothesis that the target genes are enriched in canonical biological pathways, as previously described.^49,57,58^ Supplemental data also show Core Analysis results with additional targets predicted with high confidence according to IPA, based on the TargetScan algorithm previously described.^59^ IPA designates high confidence as a cumulative weighted context score of −0.4 or lower, predicting decreased expression by at least 25% due to a specific miRNA.

The input miRNAs were reduced with higher expression thresholds so that target gene numbers did not exceed IPA software limitations for Core Analysis. The following definitions were applied for highly expressed miRNAs (baseMean>10,000 in both trimesters), similarly expressed miRNAs (P≥0.05, absolute fold-change≤2, and baseMean>1,000 in both trimesters), and differentially expressed miRNAs (FDR<0.05, absolute fold-change>2, and baseMean>1,000 in both trimesters). Due to their smaller number, no additional miRNA filters were required for Core Analysis of C14MC and C19MC targets.

### Heatmaps

Heatmap and dendrograms of samples versus miRNAs were created with a matrix of log_2_(baseMean) values scaled and centered by rows. The heatmaps and dendrograms were created with hierarchical clustering from R package *gplots* v3.1.1. Heatmaps of gene enrichment were created with R package *pheatmap* v1.0.12 with a matrix of −log_10_(P) output from IPA Core Analysis.

### Validation with qRT-PCR

Expression of 6 selected miRNAs was re-analyzed with an independent cohort by qRT-PCR using the miRCURY LNA miRNA PCR system (QIAGEN). The 6 selected (hsa-miR-144-3p, hsa-miR-24-3p, hsa-miR-126-3p, hsa-miR-145-5p, hsa-miR-143-3p, hsa-miR-126-5p) had high expression (baseMean>1000) and were significantly different in miRNA-sequencing (FDR<10^−13^) between first and third trimester. A highly expressed miRNA with stable expression in first and third trimester placentae was used as a reference gene (hsa-miR-130a-3p; baseMean>10,000 in both trimesters, P=0.9693, FDR=0.9889). RNA from first trimester (N=10) and third trimester (N=6) placenta samples were extracted, then cDNA synthesized using universal primers in the miRCURY LNA RT Kit (QIAGEN). Expression was quantified by qRT-PCR using the miRCURY LNA SYBR Green PCR Kit (QIAGEN) and a BioRad MyIQ machine, analyzed using the ΔΔCt method,^60^ with hsa-miR-130a-3p as an internal reference. Statistics were performed using the Wilcoxon rank-sum test on ΔCt values.

## Results

### Cohort demographics

There were N=113 first trimester placenta samples and N=47 third trimester placenta samples studied, including 3 subjects with both first and third trimester placenta sequenced. Principal components analysis (PCA) shows that first and third trimester placenta segregated into distinct clusters along PC1 (29.27% variability explained) and PC2 (20.22% variability explained) (Supplemental File 1). There were significantly more non-Hispanic, Caucasian parents and fetuses in the first trimester group (Table 1). However, race and ethnicity groups did not cluster in PCA analyses of the miRNA transcriptome (Supplemental File 2). Maternal pre-pregnancy BMI and thyroid disorders requiring thyroid replacement were significantly different among the groups (Table 1). There were more cases of pregnancy-induced hypertension requiring antihypertensives and/or magnesium in the third trimester placenta group, compared to none in the first trimester placenta group.

**Table 1 -.** Demographics

|  | **First Trimester** | **Third Trimester** | **P value** |
| --- | --- | --- | --- |
| **N** | 110 | 44 |  |
| **Maternal age, years** | 37.7 (3.0) | 37.3 (3.0) | 0.65 |
| **Paternal age, years** | 39.5 (4.8) | 38.6 (4.8) | 0.30 |
| **Maternal race/ethnicity** |  |  |  |
| Caucasian | 106 (96.4%) | 35 (79.6%) | 0.002 |
| Non-Hispanic | 107 (97.3%) | 35 (79.6%) | <0.001 |
| **Paternal race/ethnicity** |  |  |  |
| Caucasian | 104 (94.6%) | 37 (84.1%) | 0.04 |
| Non-Hispanic | 107 (97.3%) | 36 (81.8%) | 0.001 |
| **Fetal race/ethnicity** |  |  |  |
| Caucasian | 103 (93.6%) | 32 (72.7%) | <0.001 |
| Non-Hispanic | 106 (96.4%) | 33 (75.0%) | <0.001 |
| **Fetal female sex** | 57 (51.8%) | 18 (40.9%) | 0.22 |
| **Maternal pre-pregnancy BMI, kg/m<sup>2</sup>** | 21.9 (3.4) | 24.0 (4.7) | 0.005 |
| **Maternal pre-existing medical conditions** |  |  |  |
| Hypertension | 0 (0%) | 0 (0%) | - |
| Diabetes | 0 (0%) | 0 (0%) | - |
| Thyroid disorder | 3 (2.7%) | 6 (13.6%) | 0.02 |
| **Pregnancy complications** |  |  |  |
| Hypertension (not pre-existing) | 0 (0%) | 8 (18.2%) | <0.001 |
| Gestational diabetes | 3 (2.7%) | 1 (2.3%) | 1.0 |
| Placenta previa | 0 (0%) | 1 (2.3%) | 0.49 |
| Placental abruption | 0 (0%) | 0 (0%) | - |
| **Hypertension management in pregnancy** |  |  |  |
| Anti-Hypertensives | 0 (0%) | 3 (6.8%) | 0.022 |
| Any Magnesium use (Ante- or Postpartum) | 0 (0%) | 6 (13.6%) | <0.001 |
| **Mode of delivery- Cesarean section** | 33 (30%) | 15 (34.1%) | 0.74 |
| **Gestational age at delivery, days** | 276.3 (7.0) | 276.5 (8.0) | 0.43 |
| **Birthweight, g** | 3435.4 (463.6) | 3473.1 (467.0) | 0.7 |

Note: Values shown as mean (standard deviation) or n (%). P values were adjusted for fetal sex.

### All expressed miRNAs in placenta

We identified 2503 mature miRNAs with high-throughput sequencing of first and third trimester placentae, with 801 miRNAs reaching 10 normalized counts in both first and third trimester placentae (baseMean>10). First trimester placenta expressed 872 mature miRNAs (baseMean>10), derived from 967 miRNA precursors from all chromosomes with annotated miRNAs (22 autosomes and the X chromosome) (Figure 1A, Supplemental File 3). The majority of these precursor miRNAs originate from chromosomes 19 (12.7%), 14 (10.8%), X (9.6%), and 1 (6.8%). Third trimester placenta expressed 882 mature miRNAs (baseMean>10), derived from 985 miRNA precursors from all 22 autosomes and the X chromosome (Figure 1A, Supplemental File 3). The most represented chromosomes were also 19 (12.4%), 14 (10.5%), X (9.7%), and 1 (7.1%).

**Figure 1.**
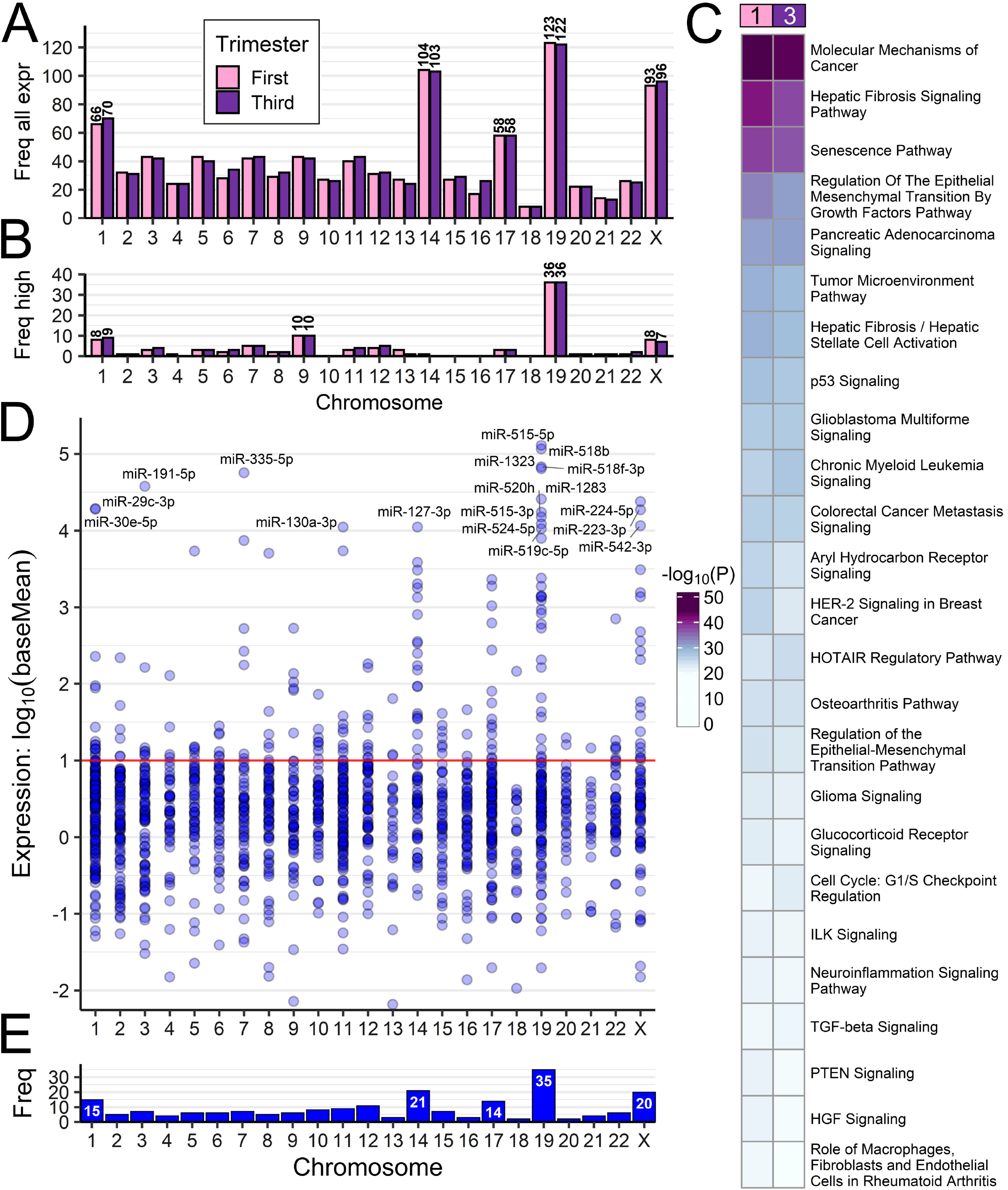
Expressed miRNAs in first and third trimester placenta. (A, B) The chromosome distribution of miRNAs expressed in placenta at first trimester (N=113 samples) and third trimester (N=47 samples). Bar plots count all genomic locations (all precursor miRNAs) corresponding to mature miRNAs identified through sequencing, at thresholds: (A) all expressed with baseMean>10 or (B) the most highly expressed with baseMean>10,000. (C) Pathway enrichment analysis with experimentally confirmed target genes of the most highly expressed miRNAs. (D) The expression distribution of miRNAs similarly expressed in first and third trimester at P≥0.05 and FC≤2. The red line (baseMean=10) is the threshold selected for stable expression. (E) Counts of similarly expressed miRNAs with P≥0.05, FC≤2, baseMean>10 in both trimesters.

### Highly expressed miRNAs in placenta

Some miRNAs had expression values several orders of magnitude higher than most miRNAs, with the median at baseMean=123.6 but the mean raised to baseMean=5,635 by these highly expressed miRNAs (Supplemental File 3). A threshold of baseMean>10,000 was selected for the “most highly expressed” miRNAs. There were 75 mature miRNAs (derived from 96 precursors) in first trimester and 77 mature miRNAs (derived from 97 precursors) in third trimester placenta which reached this threshold. The most highly expressed miRNA in first trimester was C19MC member miR-517b-3p (baseMean=218,953). The most highly expressed miRNA in third trimester and overall most highly expressed was miR-126-3p (baseMean=337,399). Chromosome 19 encoded 30 mature miRNAs (derived from 36 precursors) which reached baseMean>10,000 in both first and third trimester (Figure 1B), making chromosome 19 the source of over 37% of the most highly expressed precursor miRNAs in human placenta. Specifically, 28 of the 36 precursor miRNAs were C19MC members, and 8 localized elsewhere on chromosome 19. The next chromosomes contributing the most highly expressed miRNAs were chromosome 9, chromosome 1, and chromosome X.

We performed pathway enrichment analysis on experimentally confirmed targets of the most highly expressed miRNAs (Figure 1C, Supplemental File 5Ai). The most significantly enriched canonical pathways in first and third trimester were “Molecular Mechanisms of Cancer”, “Hepatic Fibrosis Signaling”, “Senescence”, “Regulation of the Epithelial Mesenchymal Transition by Growth Factors”, and “Pancreatic Adenocarcinoma Signaling.”

Pathway enrichment analysis with both experimentally confirmed miRNA targets as well as targets predicted with high confidence demonstrated similar patterns, though third trimester showed relatively higher enrichment in “Hepatic Fibrosis/Hepatic Stellate Cell Activation” and “Regulation of the Epithelial Mesenchymal Transition by Growth Factors” compared to first trimester (Supplemental File 4, Supplemental File 5Ai). Additional pathways, including inflammatory pathways such as “Neuroinflammation”, “Prolactin”, “Systemic Lupus Erythematosus in B Cell”, and “IL-6” signaling were also enriched (Supplemental File 5Aii).

### Similarly expressed miRNAs in first and third trimesters

There were 182 mature miRNAs with similar expression in the first and third trimester placentae (P≥0.05, fold-change≤2 and baseMean>10, Supplemental File 3), suggesting consistent expression throughout gestation (Figure 1D). These mature miRNAs are derived from 206 precursor miRNAs, with greatest representation from chromosomes 19 (17.0%), 14 (10.2%), X (9.7%), and 1 (7.3%) (Figure 1E). The most highly expressed similar miRNA was C19MC member hsa-miR-515-5p with first trimester baseMean=129,659 and third trimester baseMean=129,323, P=0.902 between trimesters. This was followed closely by other C19MC members: hsa-miR-158b, hsa-miR-518f-3p, hsa-miR-1323, and hsa-miR-1283.

### Differentially expressed miRNAs between first and third trimesters

There were 588 mature miRNAs significantly differentially expressed between first and third trimester placentae (FDR<0.05, baseMean>10) (Figure 2A), further filtered to 180 miRNAs with fold-change>2, including 91 upregulated in the first and 89 upregulated in the third trimester (Figure 2B, Supplemental File 3). The 180 differentially expressed miRNAs were derived from 202 precursors with highest representation from chromosomes 9 (9.9%), X (9.4%), 1 and 14 (8.4% each), and 19 (7.4%) (Figure 2B). The most differentially expressed miRNA was hsa-miR-4483, with 38.2-fold higher expression in the first trimester placenta (FDR=0) and a baseMean decrease from 984.1 to 25.5 from first to third trimester (Figure 2C, Supplemental File 3). The next most significantly differentially expressed miRNA was hsa-miR-139-5p with 18.1-fold higher expression in the third trimester (FDR=3.69×10^−298^), baseMean increasing from 28.0 to 497.5 (Figure 2C). The differentially expressed miRNA with the highest overall expression was hsa-miR-126-3p, with a 3.13-fold higher expression (FDR=4.52×10^−97^) in third trimester placenta (baseMean=337,399) compared to first trimester placenta (baseMean=107,787) (Figure 2D). Of the differentially expressed miRNAs, those with the greatest fold changes had lower to moderate expression around baseMean 100-1,000 and were predominantly elevated in the first trimester, whereas those with the highest expression had lower fold changes and were predominantly elevated in the third trimester (Figure 2D). Zero differentially expressed miRNAs with baseMean>1,000 reached fold-change of 4 (Figure 2D, Supplemental File 3).

**Figure 2.**
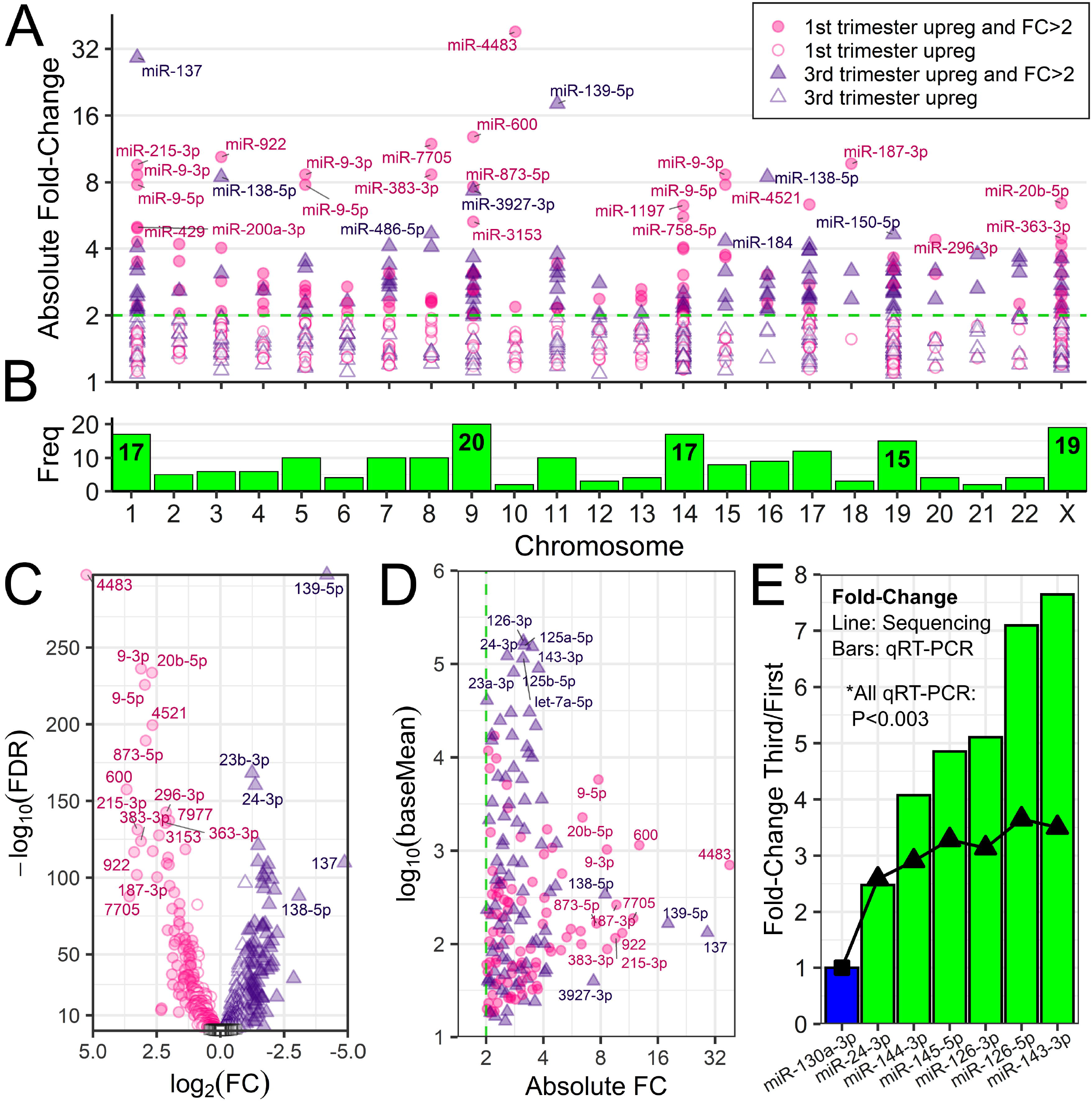
Differentially expressed miRNAs between first and third trimester placenta. (A) Scatter plot of absolute fold-change distribution across chromosomes for all differentially expressed (DE) miRNAs at FDR<0.05 and baseMean>10. The dotted line represents FC=2. (B) Chromosome frequency of 180 DE miRNA precursors at FDR<0.05, FC>2, baseMean>10. (C) Volcano plot of all miRNAs with baseMean>10. Key as in A, with addition of open black squares for non-significant miRNAs (FDR≥0.05). (D) Expression versus absolute fold-change for 180 DE miRNAs. (E) Six DE miRNAs (green) were selected for validation via qRT-PCR in an independent cohort. The bar plot shows qRT-PCR results normalized to an internal reference, hsa-miR-130a-3p (blue). The superimposed line shows fold-changes in miRNA-seq. All six miRNAs were validated significantly different between first and third trimester with P<0.003.

### Validation of differentially expressed miRNAs

Six differentially expressed miRNAs identified using NGS were selected for validation (Figure 2E). We performed qRT-PCR using an independent cohort of first (N=10) and third trimester (N=6) placenta samples. The miRNA hsa-miR-130a-3p was selected as an internal reference due to high and stable expression in first and third trimester placentae (P=0.9693, fold-change=0.9984 first/third, baseMean=11,097). All six validated miRNAs (hsa-miR-24-3p, hsa-miR-144-3p, hsa-miR-145-5p, hsa-miR-126-3p, hsa-miR-126-5p, and hsa-miR-143-3p) were upregulated in third trimester placenta with 2.5 to 3.7-fold changes by sequencing, and all six were confirmed significant by qRT-PCR with P<0.003 (Figure 2E).

### Comparison of similarly and differentially expressed miRNAs

A heatmap of the 182 similarly expressed miRNAs shows no clustering of the first and third trimester samples (Figure 3A). The heatmap of 180 differentially expressed miRNAs shows placenta sample clustering by trimester, and miRNAs clustering into two groups by direction of upregulation (Figure 3B). There was little subject variability in miRNAs in first and third trimester, but some miRNAs were not consistently expressed (baseMean=0, red).

**Figure 3.**
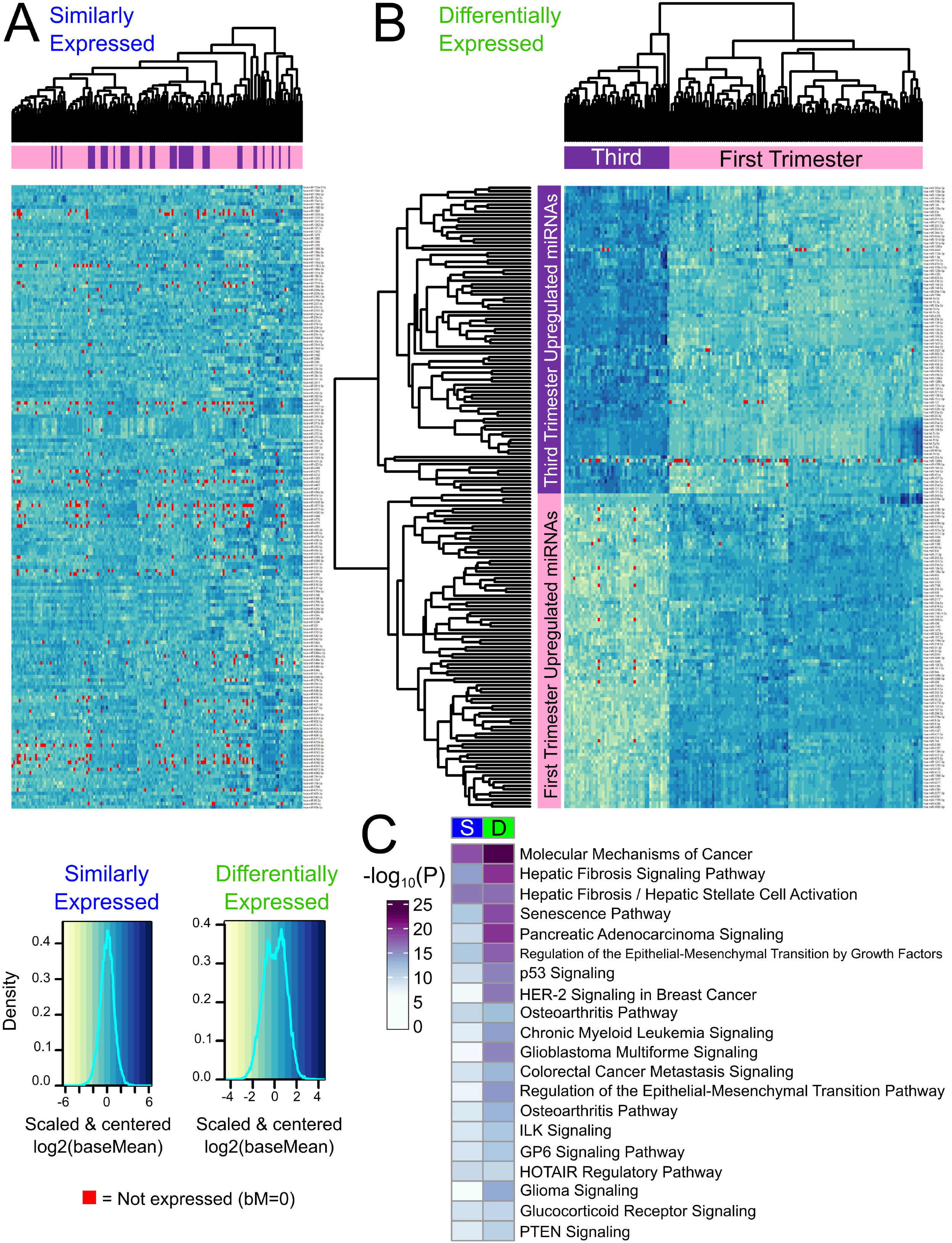
Heatmaps showing sample miRNA variability. Heatmaps: rows = scaled and centered miRNA log_2_(baseMean), columns = hierarchically clustered samples. BaseMean=0 samples are highlighted red. (A) 182 similarly expressed miRNAs with P≥0.05, FC≤2, and baseMean>10. The miRNAs are listed alphabetically. (B) 180 differentially expressed miRNAs with FDR<0.05, FC>2, and baseMean>10. The miRNAs are hierarchically clustered. (C) Pathway enrichment analysis for experimentally confirmed targets of similarly (S) and differentially (D) expressed miRNAs.

Pathway enrichment analysis was performed for experimentally confirmed miRNA targets to identify potential regulatory roles of the miRNAs expressed in placenta (Figure 3C, Supplemental File 4B). The most significantly enriched pathways, targeted by both similarly and differentially expressed miRNAs in first and third trimester placenta were “Molecular Mechanisms of Cancer” and “Hepatic Fibrosis Signaling”. None of the top 20 pathways were more significantly targeted by similarly expressed miRNAs, suggesting high variability throughout gestation (Supplemental File 5Bi). Differentially expressed miRNAs targeted more significantly by highly expressed miRNAs in the first trimester include “Molecular Mechanisms of Cancer”, “Hepatic Fibrosis Signaling”, “Senescence”, and “Regulation of the Epithelial Mesenchymal Transition by Growth Factors” pathways, suggesting these pathways are distinctly regulated by miRNAs in the first versus third trimester (Figure 3C, Supplemental File 5Bi).

When the pathway enrichment analysis was repeated with both experimentally confirmed miRNA targets as well as targets predicted with high confidence, additional patterns emerge. Differentially expressed miRNAs target the “Hepatic Fibrosis / Hepatic Stellate Cell Activation” pathway more heavily than similar miRNAs when predicted targets are included (Supplemental File 5Bii). Addition of predicted targets highlights specific cytokine and growth factor pathways, including “IL-6” and “IGF-1” signaling which are heavily targeted by similarly expressed miRNAs, and less so by differentially expressed miRNAs (Supplemental File 4, Supplemental File 5Bii).

### Expression from C14MC and C19MC

The placenta specific miRNA clusters expressed 42 mature miRNAs similarly expressed between first and third trimester placenta (P≥0.05, baseMean>1), 24 from C14MC and 18 from C19MC (Figure 4AB, Supplemental File 6). There were 105 mature miRNAs differentially expressed between first and third trimester (FDR<0.05 and baseMean>1), 64 from C14MC and 41 from C19MC (Figure 4AB, Supplemental File 6). The cluster miRNAs with highest fold-change came from C14MC: hsa-miR-1197 (6.28-fold, FDR=4.83×10^−118^), hsa-miR-758-5p (5.59-fold, FDR=5.07×10^−101^), hsa-miR-496 (4.05-fold higher in first, FDR=3.92×10^−109^), and hsa-miR-665 (3.98-fold, FDR=1.75×10^−95^), all higher in first trimester compared to third trimester placenta.

**Figure 4.**
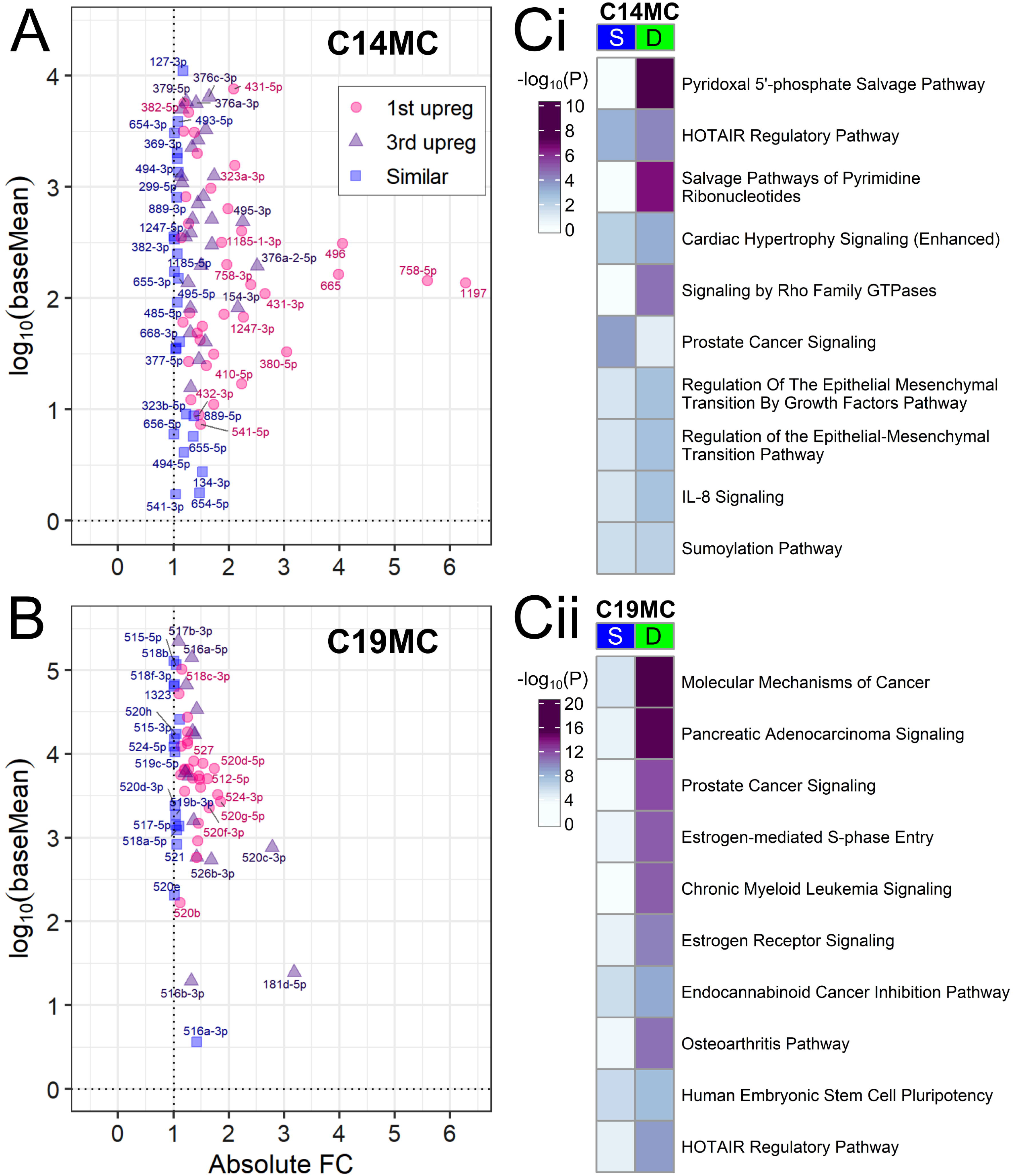
Placenta-specific C14MC and C19MC. (A,B) Expression versus absolute fold-change plots for cluster miRNAs at baseMean>1. Pink = upregulated in first trimester at FDR<0.05. Purple = upregulated in third trimester at FDR<0.05. Blue = similarly expressed with P≥0.05. Point labels are the miRNA names minus the “hsa-miR-” prefix. (A) C14MC miRNAs. (B) C19MC miRNAs. (C) Pathway enrichment analysis with experimentally confirmed targets of the S=similarly expressed or D=differentially expressed miRNAs in (i) C14MC or (ii) C19MC.

The most significantly upregulated third trimester miRNA was hsa-miR-520c-3p (2.78-fold higher, FDR=2.91×10^−111^), followed by hsa-miR-181d-5p (3.18-fold higher, FDR=6.38×10^−70^), both from C19MC. Overall, the C14 miRNA cluster contributed more differentially expressed miRNAs, reaching higher fold-changes in first trimester than C19MC (Figure 4A). Although C19MC contributed fewer total miRNAs, and at lower fold-change differences between trimesters, the C19MC baseMean distribution was an order of magnitude higher than C14MC distribution in both overall baseMean median (C19MC=5,696; C14MC=198.7) and mean (C19MC=21,278; C14MC=1,122) (Figure 4B, Supplemental File 6). This held true for both first and third trimester baseMeans expression values. Pathway enrichment analysis of experimentally confirmed target genes shows that distinct pathways are regulated by similarly and differentially expressed miRNAs (Figure 4Ci). Metabolite salvage pathways and GTPase signaling were more significantly targeted by differentially expressed C14MC miRNAs, including “Pyridoxal 5'-phosphate Salvage Pathway”, “Salvage Pathways of Pyrimidine Ribonucleotides”, and “Signaling by Rho Family GTPases” suggesting these pathways are uniquely regulated by this cluster miRNAs between the first and third trimester. “Prostate Cancer Signaling” was more significantly targeted by the similarly expressed C14MC miRNAs, suggesting this pathway is important throughout gestation (Supplemental File 5C). “Molecular Mechanisms of Cancer”, “Pancreatic Adenocarcinoma Signaling”, “Prostate Cancer Signaling”, “Estrogen-mediated S-phase Entry”, and “Chronic Myeloid Leukemia Signaling” were more significantly targeted by differentially expressed C19MC miRNAs suggesting these pathways are uniquely regulated by this cluster miRNAs between the first and third trimester. Conversely, there were no pathways among the top 30 most enriched that were targeted by similarly expressed C19MC miRNAs, suggesting more variable regulation throughout gestation (Figure 4Cii, Supplemental File D). Pathway enrichment analysis with both experimentally confirmed miRNA targets as well as targets predicted with high confidence (Supplemental File 5D), demonstrated changes in regulated pathways, with greater representation of inflammatory pathways, including the “Systemic Lupus Erythematosus in B Cell Signaling” and “Coronavirus Pathogenesis” pathways, as well as several pathways more significantly targeted by similarly expressed miRNAs, including “Role of PI3K/AKT Signaling in the Pathogenesis of Influenza”.

## Discussion

The placenta is a unique organ that changes function greatly throughout gestation, meeting different challenges and needs at different stages of pregnancy. Placentation in the first trimester sets the groundwork for its functions throughout gestation for fetal development. Placental function is in part epigenetically regulated through miRNAs, including the placenta-specific miRNA clusters, C14MC and C19MC that play critical roles in regulation of this vital organ.

This is the first study to our knowledge to use high-throughput sequencing to compare miRNA expression between first and third trimester human placentae of healthy pregnancies resulting in delivery.

The miRNA expression profiles in first and third trimester have similar chromosome distributions, with expected peaks at chromosomes 14 and 19, as well as peaks at chromosome 1, the largest human chromosome and chromosome X, which has a higher density of miRNAs compared to autosomes.^61,62^ The most highly expressed miRNA, hsa-miR-126-3p, was upregulated in third trimester, and was validated with qRT-PCR using an independent cohort. In a recent study comparing first and second trimester placenta, miR-126-3p was identified to be among the 10 most highly expressed miRNAs and identified in maternal plasma.^63^ Although variation among the first and second trimester was not different,^63^ our study identified hsa-miR-126-3p to be differentially expressed and highest in the third trimester, likely having a unique role in the third trimester compared to earlier in gestation. It is also highly abundant in fetal circulation and human umbilical vein endothelial cells,^64^ suggesting a role during parturition and fetal development and may become a potential biomarker for developmental origins of health and disease.

Among the most highly expressed miRNAs, over 37% were encoded in chromosome 19 (and 28/36 or 77.8% were specifically in C19MC), whereas none localized to chromosome 14. This supports an earlier miRNA-seq study which profiled 25 human placentae at delivery and identified higher expression from C19MC than C14MC miRNAs^65^, and additionally we show the same pattern in first trimester. The C19MC miRNAs with high expression in the placenta may potentially be used for targets, as C19MC miRNAs have been identified in maternal circulation, as early as the first trimester, with elevations throughout gestation.^66–68^ The most highly expressed miRNA that was similarly expressed was hsa-miR-515-5p, which is a member of the C19MC. Placental expression of hsa-miR-515-5p has been identified to play a key role in human trophoblast differentiation with aberrant up-regulation contributing to pathogenesis of preeclampsia.^35,69^ It has also been associated with preterm birth^70^ and fetal growth restriction.^43^ Although, it has been detected in maternal circulation, both in plasma and whole blood fractions, it has also been detected in whole blood fractions of healthy nonpregnant women,^71^ and may not be used solely as a biomarker of disease, but may be incorporated with other miRNAs with stable expression across gestation that change with disease using a bivariate biomarker disease approach described by Laurent.^45^

This atlas identified 180 differentially expressed miRNAs which may be important for functional changes in the placenta throughout pregnancy. Among those, the most differentially expressed miRNAs were highest in the first trimester. The most significantly targeted pathways of experimentally confirmed targets by differentially expressed miRNAs and the most highly expressed in the first trimester was “Molecular Mechanisms of Cancer.” Although identified in pathway enrichment analyses for tumor progression, many of the major signaling pathways involved in inter- and intra-cellular communication of invasive phenotypes mimic those associated with migration and invasion of trophoblasts into the maternal decidua and spiral arteries. These essential placentation steps take place in an environment rich in hormones, cytokines, and growth factors and include responsible signaling pathways such as mitogen-activated protein kinase (MAPK), phosphoinositide 3-kinase (PI3K)/protein kinase B (Akt), Janus kinase (JAK)/signal transducer and activator of transcription proteins (STATs), wingless (Wnt), and focal adhesion kinase (FAK) pathways.^2^ Of these miRNAs, the most differentially expressed with highest expression in first trimester, hsa-miR-4483, was also found to be strongly downregulated in the second trimester and hence likely plays a significant role in the first trimester placenta. It may function to regulate estradiol production early in gestation, as described in another hormone producing cell type and contribute to migration and invasion.^72^ Differentiation of first trimester human placental cytotrophoblasts from an anchorage dependent epithelial phenotype into the mesenchymal-like invasive extravillous trophoblast is a crucial step for placentation. Illsley et al, previously demonstrated that an epithelial to mesenchymal transition takes place when first trimester cytotrophoblasts differentiate into extravillous trophoblasts.^73,74^ MiRNAs have been implicated in the epithelial to mesenchymal transition.^75–80^ The most highly differentially expressed miRNA in first trimester placenta, hsa-miR□205 has been implicated in the epithelial to mesenchymal transition and the maintenance of the epithelial phenotype.^75–79^ In human trophoblast cell lines it has been identified to silence MED1 under hypoxic conditions.^75^ suggesting it has a role in the first trimester regulating trophoblast differentiation during physiologic hypoxic conditions.^75,81,82^ “Regulation of the Epithelial Mesenchymal Transition by Growth Factors Pathway” was also one of the most significantly targeted pathway by differentially expressed miRNAs and most highly expressed in the first trimester, highlighting the differences between placentation when the placenta is invading maternal tissue and establishing itself in states of low oxygen tension versus time of delivery when the placenta has completed its purpose.

Two additional significantly targeted pathways by differentially expressed miRNAs and highly expressed in the first trimester included the “Hepatic Fibrosis Signaling” and the “Senescence” pathways. Hepatic Fibrosis Signaling is classically associated with extracellular matrix deposition,^83^ consistent with first trimester placental function when extravillous trophoblasts degrade and induce ECM remodeling to enable migration.^2,5,84,85^ Cellular senescence is programmed cell-cycle arrest that restricts the propagation of cells, which is induced by various forms of cellular stress, including oxidative stress. Cell fusion, has also been identified to trigger cellular senescence and has been described in the placenta, with the placental expressed fusogen, Syncitin-1 (*ERVWE1)*, which mediates cell-fusion-induced senescence of the syncitiotrophoblast.^86–88^ These senescent cells secrete inflammatory cytokines, chemokines and matrix metalloproteinases, known as the senescence associated secretory phenotype (SASP). SASP proteins promote EMT and the degradation of basement membranes, increasing migration and invasion for appropriate placentation.^89^

Our findings also support the importance of the placenta-specific miRNA clusters throughout gestation, with 42 miRNAs similarly expressed and 105 differentially expressed across first and third trimester. This indicates that while they are placenta-specific miRNAs, the majority have varying roles throughout pregnancy. Differentially expressed canonical pathways targeted by the C14 and C19 clusters were more significant than those of all similarly expressed miRNAs suggesting these clusters have significantly different roles throughout gestation.^66^ Similar to other studies using whole villous tissue and primary cytotrophoblasts,^20,48,63^ we identified a decrease in C14MC expression from first to third trimester. However, a recent study did not identify a decrease throughout gestation, but their study only focused on the first and second trimester of presumably normal pregnancies.^63^

The major strengths of this study are the use of first and third trimester tissue from healthy pregnancies resulting in delivery, the cohort size, the availability of detailed demographic information, and the use of high-throughput sequencing. NGS, as opposed to other techniques such as array, allows for greater confidence in the conclusions regarding differential expression, since all known miRNA species previously annotated in the human genome are considered, and bias is not introduced by eliminating certain RNAs. Previous studies analyzing miRNA expression in first and/or third trimester placentae have used microarray technology and most examined very few samples (N=2-6 in each group).^25,47,48,90^ There are currently few NGS miRNA profiles of the placenta, and our study is the first to profile both first and third trimester placentae with NGS and a large sample size. We successfully validated all six selected miRNAs using qRT-PCR and an independent cohort.

Our study has some limitations. There were some differences in the demographics between the groups from the first and third trimester placenta samples. This includes race, ethnicity, maternal BMI, thyroid disorders, and pregnancy complications, specifically hypertension. However, the overall differences were small. In addition, PCA analysis did not demonstrate outliers. Furthermore, we performed pathway enrichment analysis using only experimentally confirmed targets. When performed using both experimentally confirmed and predicted with high confidence targets, although overall pathways and patterns remained consistent, when we only included experimentally confirmed targets, immune mediated pathways were not represented. Overall, we intended to identify and compare the normative miRNA signatures in the first and third trimester placentae. Our study shows many stably expressed miRNAs throughout gestation as well as significant differences between the miRNA signatures. This work provides a rich atlas to direct functional studies investigating the epigenetic differences in first and third trimester placentae and development of disease related biomarkers or prognostic indicators that are gestational age specific.

## Future Perspective

As we improve our understanding of miRNA profiles in placenta and across gestation, miRNAs may be useful biomarkers for non-invasive prenatal diagnostic testing. Our knowledge of miRNA profiles is still in its infancy relative to our knowledge of the protein coding transcriptome. Until recently, most miRNA profiling papers of placenta used arrays with limited samples. However, protocols to capture small RNAs, synthesize cDNA, and perform high-throughput NGS are improving rapidly. In 5-10 years’ time, we expect that the knowledge of human miRNA profiles in different tissue and extracellular locations will greatly improve as well. This will provide opportunities for biomarker discovery and diagnostic test development, since miRNAs are smaller, more stable RNAs than protein coding transcripts. Currently, the knowledge pool of miRNA targets has limited confirmed miRNA-RNA interactions, but this will improve as the miRNA field continues to evolve. Our work to profile miRNAs in first and third trimester provides a foundation for biomarker discovery during pregnancy and future advancements in maternal-fetal health.

## Executive Summary

- This work creates an atlas of the miRNA expression profiles of first and third trimester human placenta from patients who delivered healthy babies.
- Chromosome 19 contributes approximately 37% of the most highly expressed miRNAs in both first and third trimester placenta. Most of these miRNAs are localized to the pregnancy-associated miRNA cluster, C19MC.
- There are 182 miRNAs with similar expression across gestation. Other patient variables may affect the abundance of these miRNAs.
- There are 180 miRNAs with significant differences in expression between first and third trimester placenta. These miRNAs may contribute to changes in placental function or be markers of different placental stresses throughout gestation.
- Six miRNAs were successfully validated with qRT-PCR in an independent cohort.
- The placenta-specific miRNA clusters (C14MC and C19MC) contain both similarly and differentially expressed miRNAs.
- C14MC expressed miRNAs with greater fold-change differences across gestation than C19MC miRNAs, though C14MC miRNAs are not among the most highly expressed miRNAs in placenta.
- For both similarly and differentially expressed miRNAs, C19MC miRNA placenta expression was overall higher than C14MC expression.

## Supplemental Information

**Supplemental File 1. Principal Components Analysis, fetal sex**. PCA plot for the miRNA-seq results of N=113 first trimester and N=47 third trimester placenta samples. The samples cluster by trimester. Samples are color-coded by trimester group and fetal sex, female (F) and male (M). Three patients with matched samples are labeled. [PDF]

**Supplemental File 2. Principal Components Analysis, race and ethnicity.** PCA plots for the miRNA-seq results of N=113 first trimester (CVS) and N=47 third trimester (PL) placenta samples. Samples shapes indicate race and ethnicity. [PDF]

**Supplemental File 3. Analysis of differential miRNA expression between first and third trimester human placentae.** Tables of mature miRNAs DESeq2 results annotated with precursor and chromosome information. (A) Mature miRNAs with no duplicate rows. (B) Rows split by the chromosome column, for scatter plots. (C) Rows split by both the chromosome and precursor miRNA columns, for bar plots to count all genomic locations. (D) DESeq2 results with normalized counts for each sample. [Excel .xlsx]

**Supplemental File 4.**Compilation of full target gene enrichment analysis results from IPA Core Analysis. [Excel .xlsx]

**Supplemental File 5.** Extended pathway enrichment analysis heatmaps for (i) only experimentally confirmed miRNA targets or (ii) both experimentally confirmed and high confidence predicted miRNA targets. (A) Highly expressed miRNAs with baseMean>10,000 in first (pink, “1”) or third (purple, “3”) trimester. (B, C, D) Similarly (blue, “S”) and differentially (green, “D”) expressed miRNAs encoded by (B) all chromosomes, (C) C14MC, (D) C19MC. Heatmap data are −log_10_(P) output from IPA Core Analysis. [PDF]

**Supplemental File 6.** Subset of Supplemental File 3 spreadsheets with placenta-specific miRNA clusters, C14MC and C19MC. [Excel .xlsx]

