## Supplementary figures and images for "High-throughput miRNA-sequencing of the human placenta: expression throughout gestation"

### Supplemental File 1

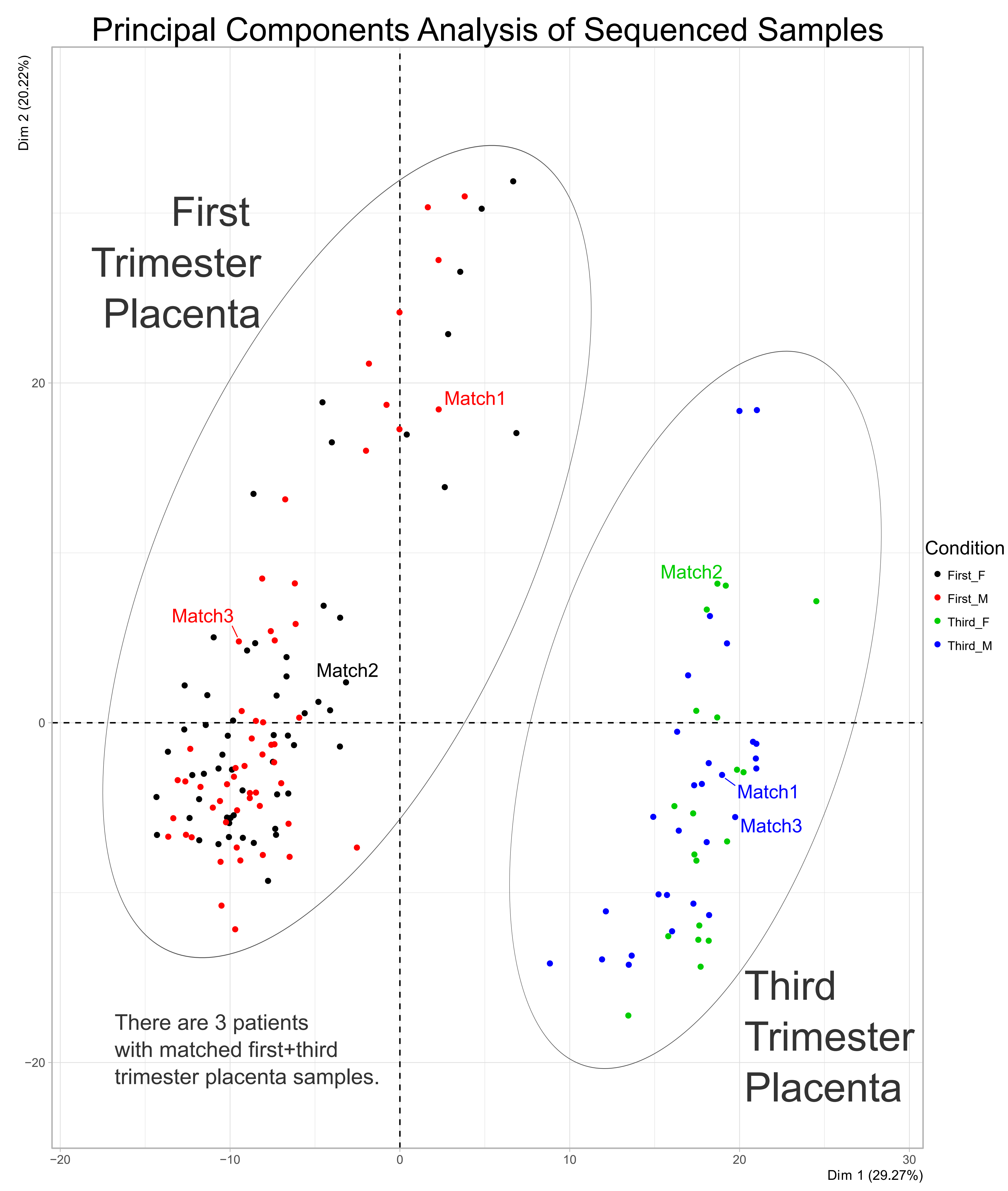

### Supplemental File 2

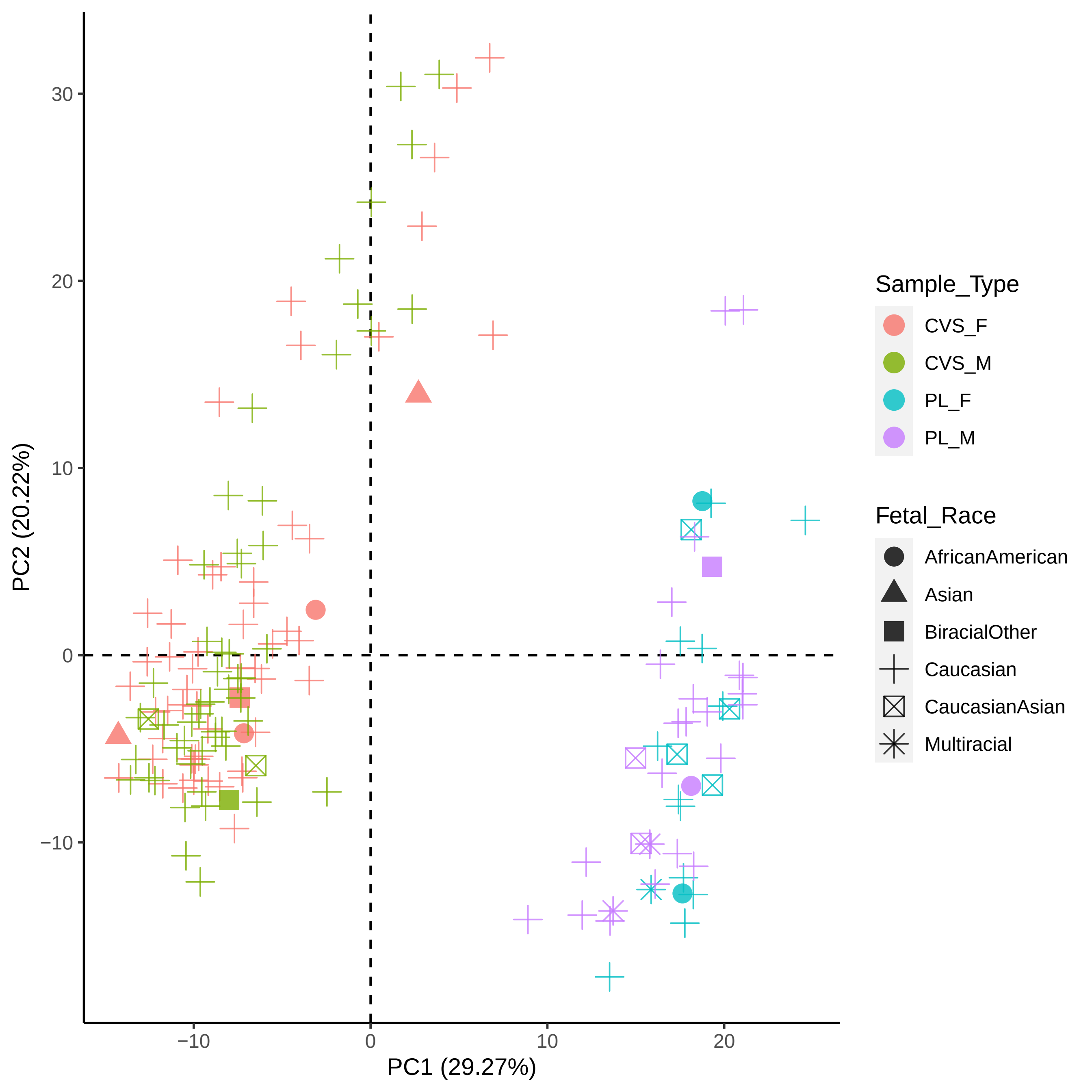

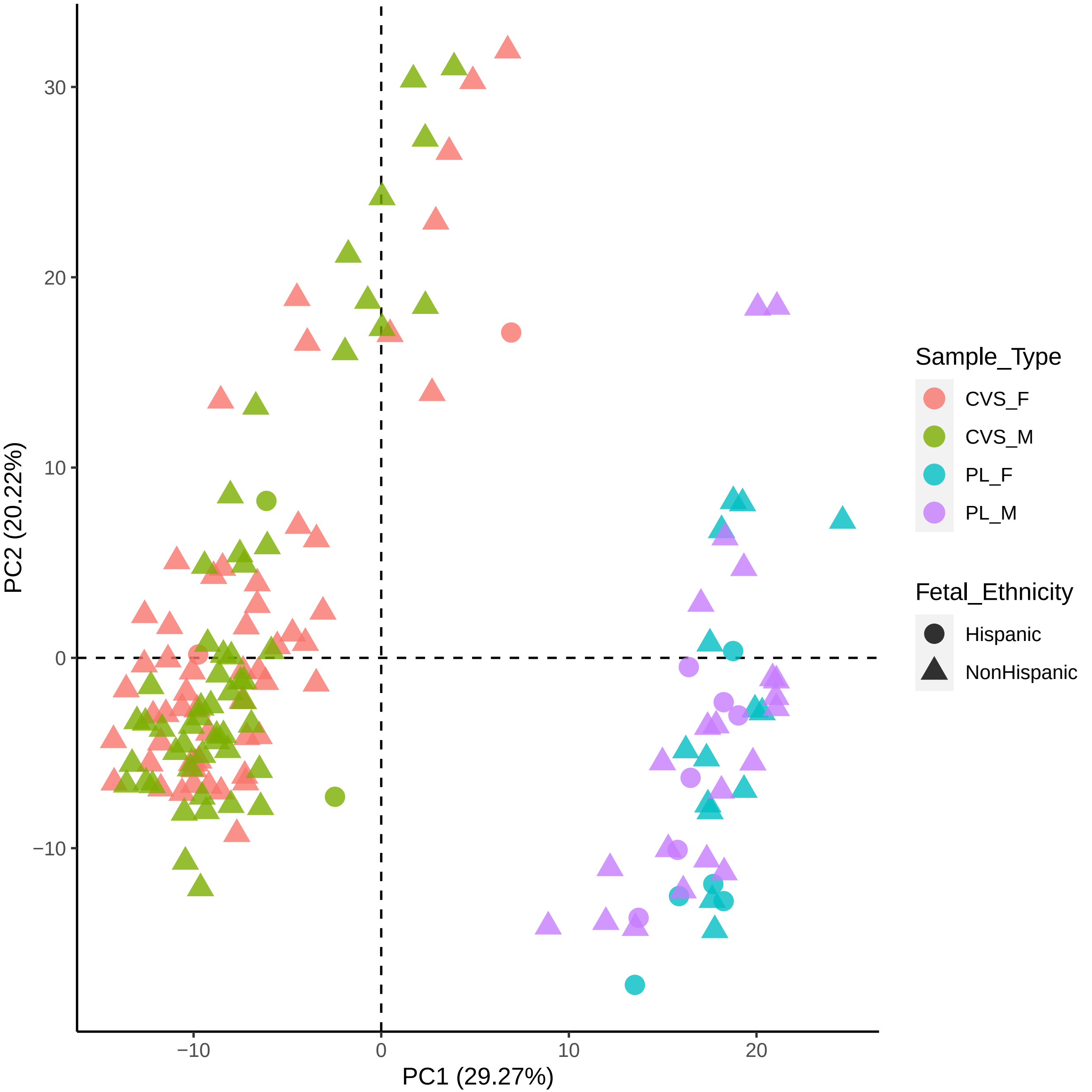
