## Supplemental File 5 for "High-throughput miRNA-sequencing of the human placenta: expression throughout gestation"

### All Highly Expressed miRNAs

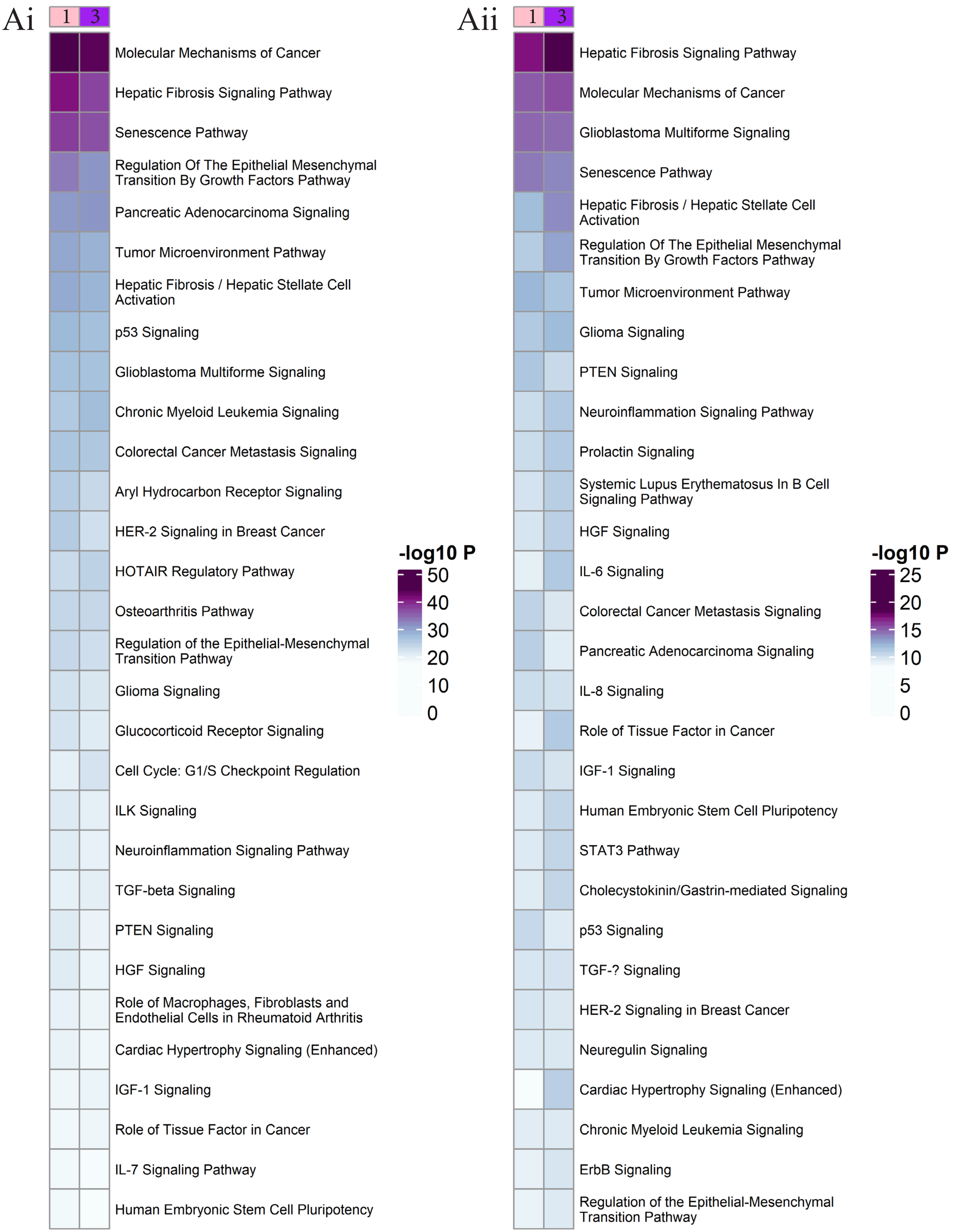

### Similarly and Differentially Expressed miRNAs

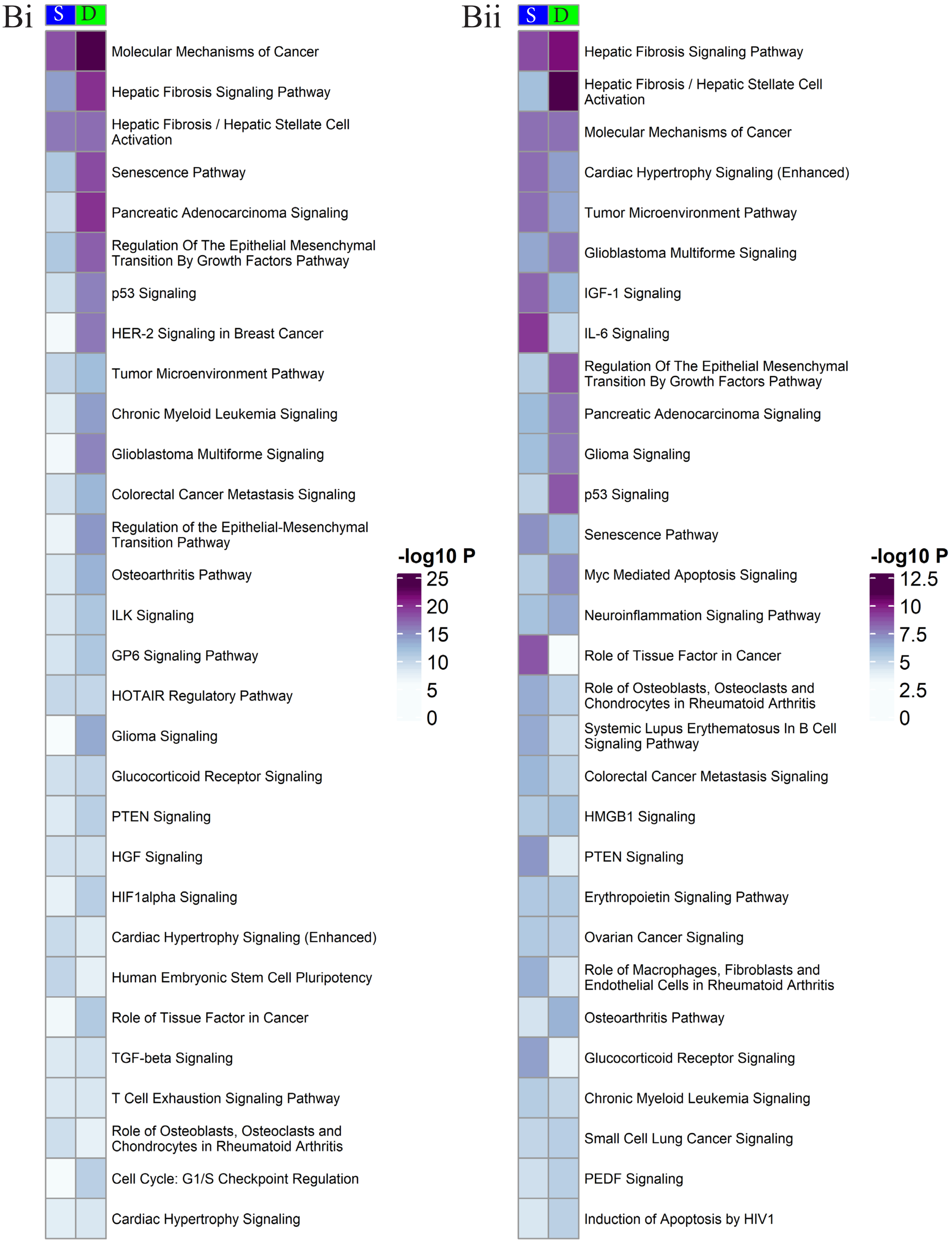

### C14MC Expressed miRNAs

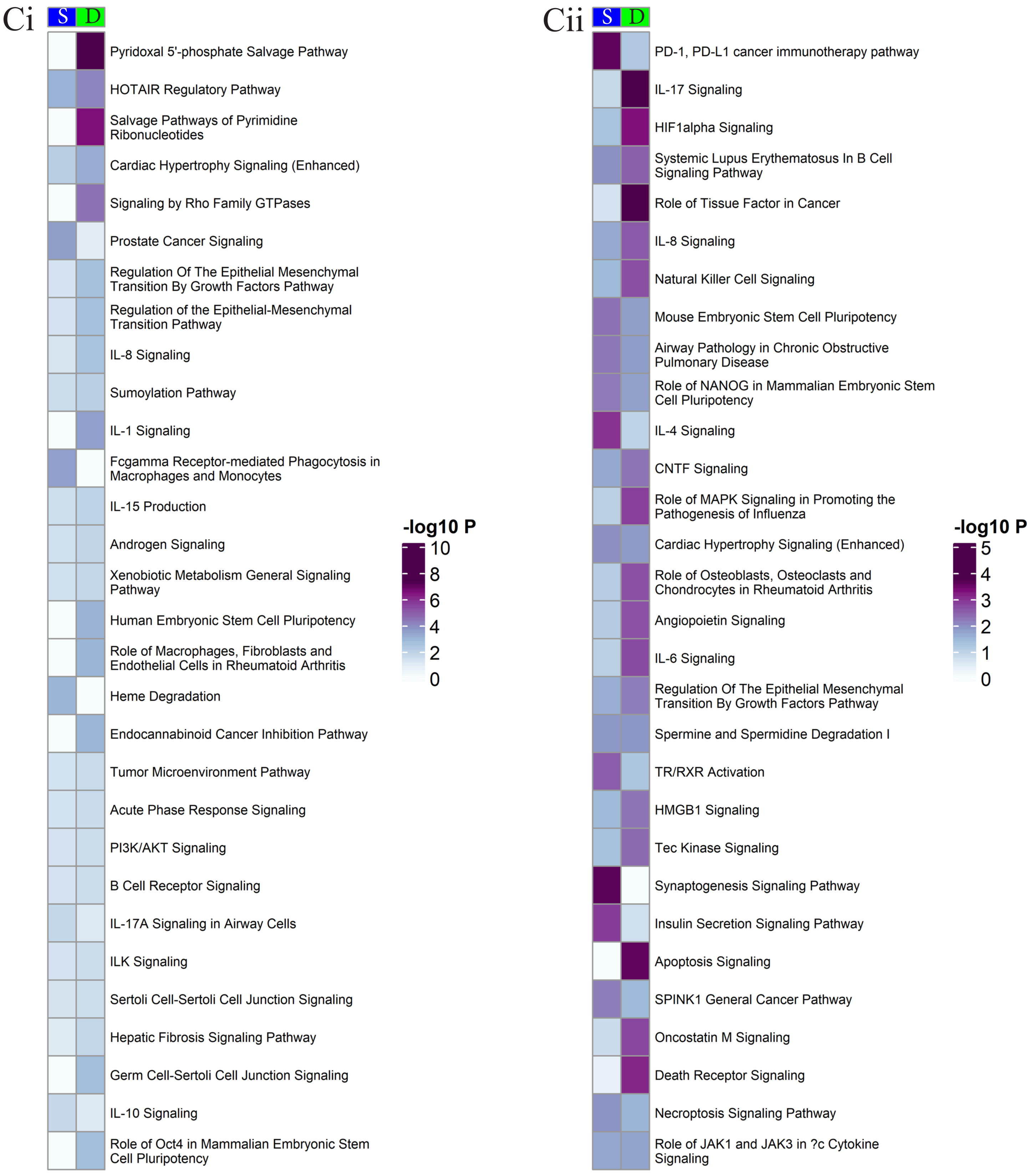

### C19MC Expressed MiRNAs

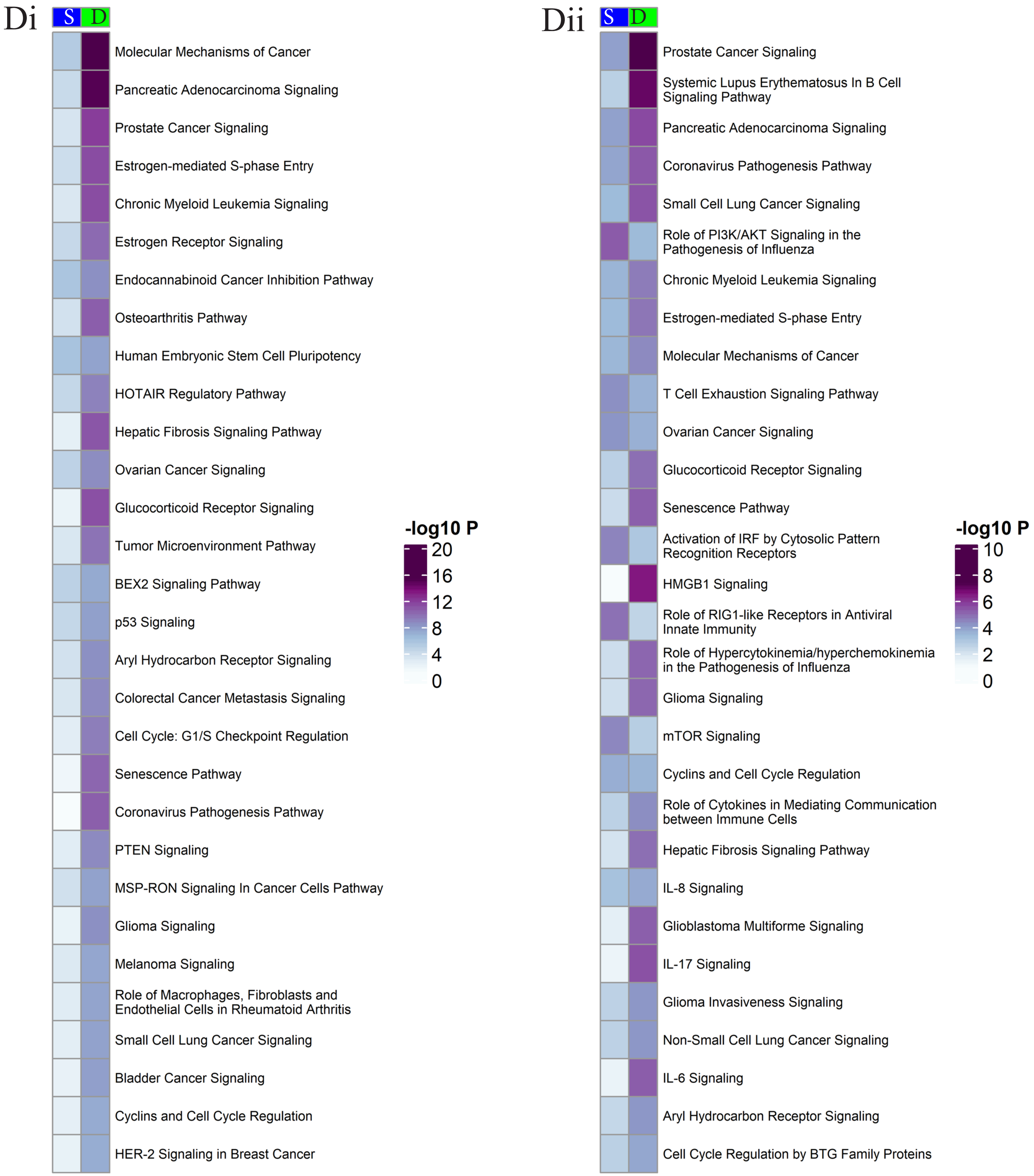
